# Deep origins, distinct adaptations and species level status indicated for a glacial relict seal

**DOI:** 10.1101/2025.02.06.636115

**Authors:** Ari Löytynoja, Jaakko Pohjoismäki, Mia Valtonen, Juha Laakkonen, Wataru Morita, Mervi Kunnasranta, Risto Väinölä, Morten Tange Olsen, Petri Auvinen, Jukka Jernvall

## Abstract

Isolated populations of postglacial relicts are known from many regions and are typically found on mountains for terrestrial species and in lakes for aquatic species. Among the few aquatic mammalian relicts, the Saimaa ringed seal (*Pusa hispida saimensis*) has been landlocked in Lake Saimaa, Finland, for the last 10,000 years. Saimaa ringed seals show genetic, behavioral, and morphological differences to the other ringed seal subspecies, but the extent these differences stem from the end of the last glacial period remains unclear. Here, we demonstrate with comprehensive sampling and state-of-the-art genomic methods that the Saimaa ringed seals are much older than the lake they inhabit, having formed a separate evolutionary branch for at least 60,000 years. This deep evolutionary origin of the Saimaa ringed seals is further underscored by our ecomorphological analyses revealing adaptively distinct features in their dentition and tongue. Overall, glacial relicts, many threatened by extinction, may harbor a richer selection of evolutionary history than might be expected from their recent isolation history alone.

## Introduction

Ringed seals *Pusa hispida* (Schreber, 1775) are the most common Arctic pinnipeds, found throughout all seasonally ice-covered northern seas and two freshwater lakes (Figure 1A). They are highly ice-associated, adapted to maintaining breathing holes and constructing subnivean lairs for parturition, nursing, and resting (McLaren 1958, Kelly 2022). The nominate subspecies, *P. hispida hispida* (Schreber, 1775) (hereafter the “Arctic ringed seal”), is the most widespread, inhabiting the circumpolar Arctic Ocean. Three subspecies are present as isolated populations in the Fennoscandia (Figure 1B), the Baltic ringed seal *P. h. botnica* (Gmelin, 1788), the Ladoga ringed seal *P. h. ladogensis* (Nordqvist, 1899), and the Saimaa ringed seal *P. h. saimensis* (Nordqvist, 1899). The presence of these three Fennoscandian subspecies has been explained by the isolation of the ringed seals into the Baltic Sea basin after the last glacial period with further entrapment of seals into Lake Ladoga and Lake Saimaa (Ukkonen et al. 2014). Lake Saimaa is a freshwater lake with a surface area of 4400 km^2^ and houses a population of close to 500 endemic ringed seals, making these one of the most endangered pinnipeds in the world (Kovacs et al. 2012, https://www.metsa.fi/en/nature-and-heritage/species/saimaa-ringed-seal/). Currently, the population is slowly growing due to active conservation efforts, but the Saimaa ringed seal is still threatened by bycatch mortality and climate change driven habitat change (Kunnasranta et al. 2021).

**Figure 1.**
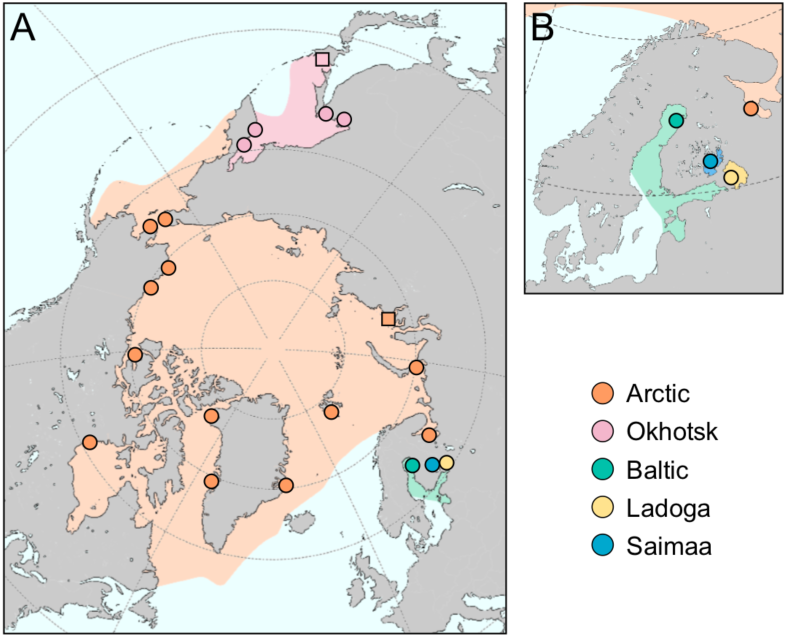
The distribution and sampling of the studied ringed seals and their distribution ranges. (A) The nominal subspecies of the Arctic ringed seal (*Pusa hispida hispida*) has circumpolar distribution while the four other subspecies are geographically restricted. (B) The ringed seals living in the Fennoscandia are made of landlocked populations in the Lake Saimaa and Lake Ladoga, and the Baltic Sea population. Phenotypic sampling from museum collections roughly matches those of the genetic data, except for Arctic samples from Kara Sea (square) substituting for Pechora Sea, and Okhotsk samples being from Hokkaido or nearby islands (square).

The morphological and ecological differences between the Saimaa ringed seal and the other subspecies have been long known (Nordqvist 1899, Sipilä et al. 1990, Hyvärinen and Nieminen 1990, Sipilä and Hyvärinen 1998, Kunnasranta 2001, Amano et al. 2002), historically assumed to derive from the Holocene postglacial isolation of Saimaa population from Baltic Sea population. However, genetic analyses of mitochondrial DNA (mtDNA) haplotypes (Valtonen et al. 2012, 2014, Heino et al. 2023), microsatellite loci (Palo 2003, Palo et al. 2003, Nyman et al. 2014), and genome-wide variation (Löytynoja et al. 2023, Sundell et al. 2023) have revealed greater genetic differences between Saimaa and other populations than can be explained by postglacial drift and recent adaptation.

In this study, we explore more broadly the genomic differences between the Saimaa and other ringed seals by including the previously unsampled Arctic ringed seals in northern Eurasia, as well as the Okhotsk ringed seal *P. h. ochotensis* (Pallas, 1811), an Asian Pacific representative of the species (Figure 1). In addition to mitochondrial and nuclear genomes, we examine feeding morphology differences to address the magnitude of ecological differentiation among the subspecies. We show that even with circumpolar sampling of Arctic ringed seal populations, the Saimaa ringed seal retains its genetic uniqueness. Based on the genomic divergence from the other ringed seals, together with specialized morphological features linked to feeding ecology, we argue that the Saimaa ringed seal is evolutionary more unique than previously thought.

## Results

### Genetic origins and differentiation of the Saimaa ringed seal

Our phylogenetic analysis of mtDNA data, representing the first truly global sampling of mitochondrial diversity in ringed seals, confirms earlier findings and shows Saimaa forming a tight cluster with closely-related haplotypes present but rare among the Baltic and Arctic ringed seals (Figure S1) (Valtonen et al. 2012, Nyman et al. 2014, Heino et al. 2023, Olsen et al. 2025). The wider sampling from the eastern hemisphere fails to establish connections between Saimaa and other populations. Similarly to Ladoga and Baltic ringed seals, the haplotypes of the sampled Okhotsk individuals are spread into multiple distinct lineages of the phylogenetic tree (Figure S1). In contrast, despite being embedded among the global mtDNA variation, the Saimaa population appears to form the only monophyletic lineage of mtDNA haplotypes.

Compared to the nonrecombining mtDNA haplotypes, the nuclear genome is expected to provide a more nuanced view into population histories. Still, the Saimaa sample forms a tight cluster in the principal component analysis (PCA) based on 1.808 million SNPs across the whole nuclear genomes from five individuals per population. The Saimaa sample is separated from all the others, including the Okhotsk ringed seal, by PC1 whereas the other populations are separated by PC2 (Figure 2A). The full 46 individual dataset produces largely similar division between the populations but places the Arctic and Okhotsk samples closer to each other (Figure 2B), possibly as an artifact of uneven sampling (Elhaik 2022). Furthermore, the derived allele statistics show that the Saimaa ringed seals retain a high representation of population-specific variants, a pattern preserved when multiple individuals are compared (Figure 2C). The relative magnitude of the private variants in Saimaa makes it highly implausible that this variation has resulted from drift during the 10 kya, or roughly 1000-generation entrapment in the lake. This raises the question how deep are the roots of the Saimaa population.

**Figure 2.**
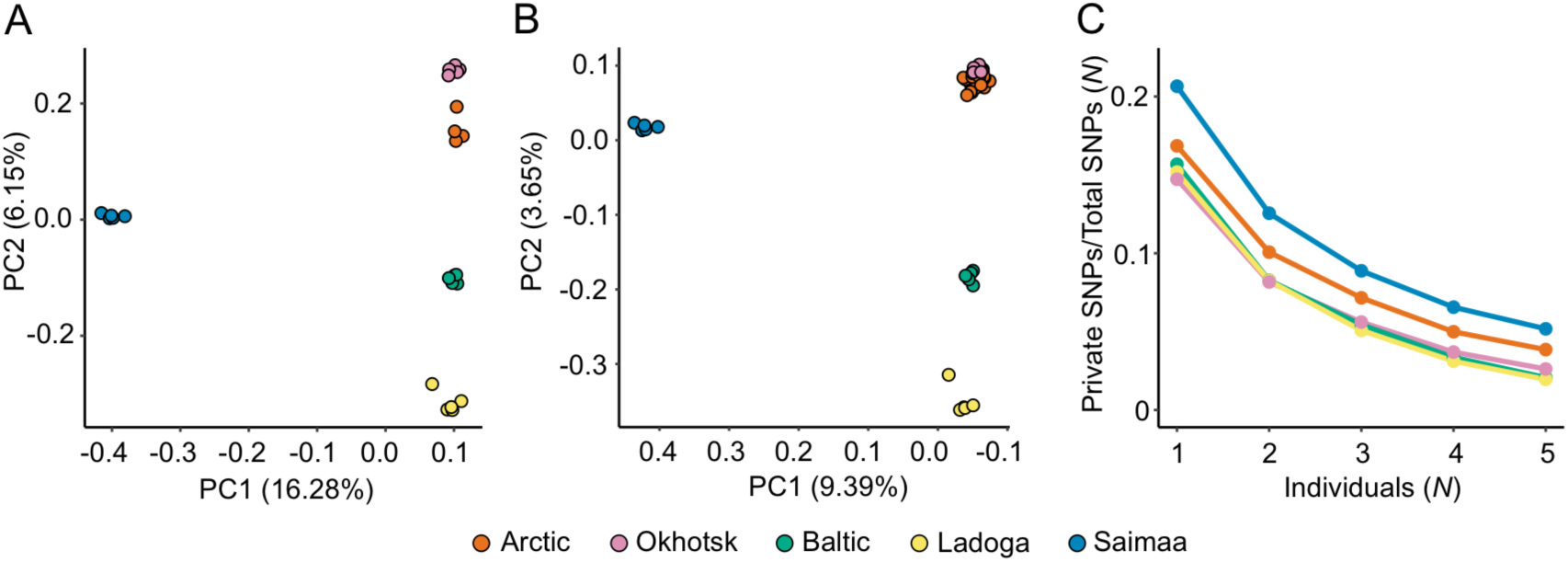
Genetic uniqueness of the Saimaa ringed seals. (A) Using five individuals from each population, the Saimaa ringed seals form a tightly defined and separate cluster along the PC1 in the PCA analysis, while the genetic differentiation of the non-Saimaa seals creates a geographic continuum along the PC2: the Baltic are placed between the Ladoga and Arctic, and Okhotsk further away. The Arctic individual placed between the rest of the Arctic and Okhotsk is from the Bering Strait. (B) Comparable patterns are obtained using the full 46 individual data, although the uneven sampling hides the separation between Arctic and Okhotsk. The x-axis is reversed to visually match (A). (C) With MAF 0.05 filtering, Saimaa has the highest proportion of population-specific derived alleles in samples of 1-5 individuals per subspecies.

To better understand the population histories, we focused on the divergence times among the three Fennoscandian populations (Baltic, Ladoga, Saimaa) in reference to the Arctic population using MSMC-IM (Wang et al. 2020), a demographic inference method providing time-dependent estimates of population-specific *N_e_* as well as gene flow. Even with this improved modeling compared to a previous analysis (Löytynoja et al. 2023), the inferred *N_e_* trajectories for the Saimaa population remain distinctly different from those of other populations throughout most of the late-glacial period. Notably, the Saimaa population shows a dramatic decline that appears to begin before the seals could enter the present lake basin (Figure 3A). In contrast, the other populations follow similar trajectories in the deeper time, prior to their postulated split. The estimates of migration rate over time (Figure 3B) demonstrate the recent strict isolation of the Saimaa population but also suggest the Saimaa lineage to have been in contact with the Baltic and Ladoga population around 8–10 kya and the exchange been at its greatest around 50–80 kya (Figure 3B). The gene flow among the three Fennoscandian populations between 8–10 kya provides an explanation for the presence of Saimaa-associated mtDNA haplotypes and microsatellite alleles – although at low frequency –in the Baltic population (Palo 2003, Palo et al. 2003, Valtonen et al. 2012, Nyman et al. 2014). On the other hand, compared to the Baltic, the slightly earlier isolation of Ladoga ringed seals from the Arctic (Figure 3B) can explain the reduced mtDNA haplotype variation observed in that population (Figure S1).

**Figure 3.**
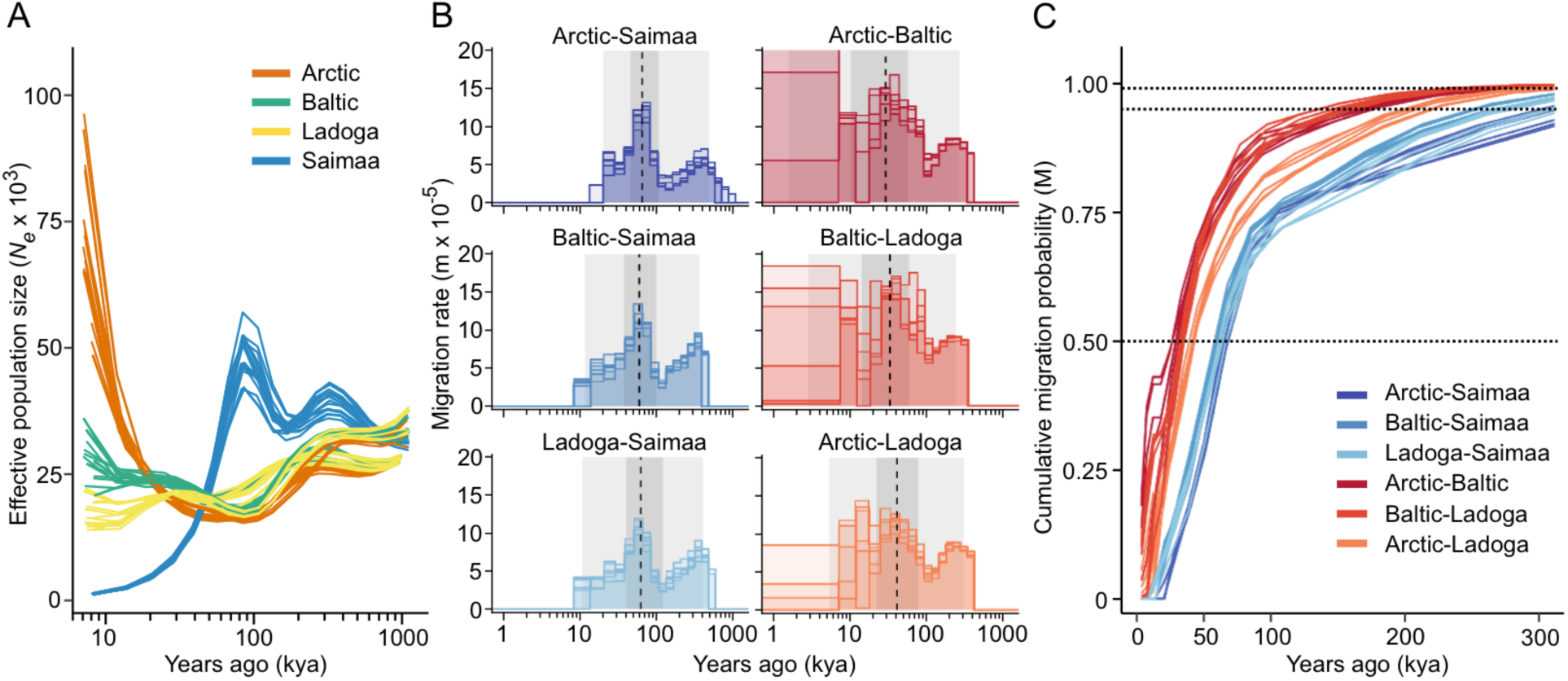
Estimated past effective population sizes and migration rates reveal a deep history for Saimaa ringed seals. (A) Estimates of past effective population sizes (*N_e_*) for the three Fennoscandian and Arctic ringed seals (the distant Okhotsk is not included in these analyses) as computed from the full genome data. (B) Estimates of migration rates over time between pairs of individuals from different populations. The rates are cut at the point where the cumulative migration probability reaches 0.999. Background shadings indicate the average cumulative probability of 0.01, 0.25, 0.75 and 0.99, and dashed lines that of 0.50. (C) Cumulative migration probability between pairs of individuals from different populations. The dotted lines indicate the thresholds M_50_, M_95_ and M_99_, the x-coordinates of intersections defining the age of each pair reaching the respective level. Five chromosomes per population were included and each line represents a comparison of individuals from two different populations. Mutation rate (!) and generation time were 1.826e-8 and 10 years, respectively.

To further date the roots of the Saimaa lineage, we computed M, the cumulative migration probability over time, between each population pair (Figure 3C). These confirm the clearly distinct accumulation of gene flow between Saimaa and the three other populations, while the separate trajectories for the Arctic-Ladoga pairs support multiple waves of gene flow from the Atlantic to the Baltic region, some of that not reaching Ladoga. As a proxy for the population split times (Wang et al. 2020), we estimated the times for M reaching 50% (M_50_) as well as the more stringent thresholds of 95% (M_95_) and 99% (M_99_) (Table 1). With the applied parameters of mutation rate (*µ*) and generation time (*g*), M_50_ for Saimaa varies between 60.2 kya (with Baltic) and 65.8 kya (with Arctic). These values are about twice as high as those among the three non-Saimaa populations, varying from 29.2 kya to 41.7 kya (Table 1). In comparisons including Saimaa individuals, the average M_95_ are over 261 kya, indicating that parts of the Saimaa genome have even deeper ancestry (Table 1). It is also notable that the M cut-off values for the Saimaa-Ladoga pairs are deeper in the past than they are for the Saimaa-Baltic pairs, highlighting the distinct origins of the two lake populations.

**Table 1.** Average age (years ago) for cumulative migration probability reaching 50%, 95% and 99% for different population pairs. SD=standard deviation.

|  | M<sub>50</sub> | SD<sub>50</sub> | M<sub>95</sub> | SD<sub>95</sub> | M<sub>99</sub> | SD<sub>99</sub> |
| --- | --- | --- | --- | --- | --- | --- |
| Arctic-Saimaa | 65,819 | 1,770 | 336,303 | 29,985 | 490,069 | 50,029 |
| Baltic-Saimaa | 60,220 | 1,692 | 261,615 | 10,421 | 372,045 | 14,322 |
| Ladoga-Saimaa | 63,152 | 2,033 | 294,702 | 21,567 | 413,519 | 32,510 |
| Arctic-Baltic | 29,199 | 2,874 | 169,737 | 5,719 | 272,638 | 6,849 |
| Baltic-Ladoga | 33,502 | 1,151 | 148,504 | 10,678 | 244,162 | 12,628 |
| Arctic-Ladoga | 41,718 | 1,369 | 209,608 | 8,912 | 310,000 | 14,762 |

The early isolation of the Saimaa lineage between 10–60 kya could be explained by a refugium in the glacial lakes either on the eastern or southern side of the Fennoscandian ice sheet. During 90–50 kya, large glacial lakes existed in the West Siberian Plains and White Sea Basin with water routes to the eastern edge of the Fennoscandian ice sheet (Mangerud et al. 2001, Mangerud, et al. 2004, Krinner et al. 2004). Similarly, a chain of ice lakes existed at the southeastern border of the glacier, in current-day western Russia and Belarus, with connections to the Caspian Sea and Black Sea (Gorlach et al. 2017). After the latest glacial maximum, around 11.6 kya, the Baltic Ice Lake drained to the Atlantic, creating a brief phase of brackish-water Yoldia Sea (Patton et al. 2017) followed by the re-enclosed Ancylus Lake (Bjorck 1995). Migration of seals between the Atlantic and the Baltic Sea basin has been permanently possible since the opening of the connection through the Danish Straits 8.5 kya (Ukkonen et al. 2014), and also detectable in the genetic data (Martinez-Bakker et al. 2013).

Taken together, the genomic evidence demonstrates that Saimaa ringed seals represent an evolutionary lineage with deep roots, clearly distinct from the contemporary Baltic and Ladoga ringed seals that were post-glacially derived from the oceanic population. The approximately two-fold divergence time indicates that the ancestors of the Saimaa population might have become isolated in an ice-dammed lake system east or southeast of the continental ice sheet already during the build-up of the last Fennoscandian Ice sheet. Regardless of the precise geographic scenario, and used mutation rate and generation time parameters (see Methods), the population size trajectory points to an even older independent history for the Saimaa lineage (Figure 3). Along with this unique ancestry, the Saimaa lineage also carries ancestry derived from a postglacial admixture with Atlantic ringed seals. Currently, the land-locked Saimaa ringed seals are effectively allopatric and thus continuing their distinct evolutionary trajectory into the foreseeable future.

### Phenotypic differentiation of the Saimaa ringed seal relating to feeding ecology

Given the deep ancestry of the Saimaa lineage, next we examined phenotypic features that may relate to feeding ecology differences between marine and freshwater habitats. Previous analyses have demonstrated morphological features, especially related to the skull that can be used to differentiate Saimaa ringed seals from other ringed seals (Nordqvist 1899, Hyvärinen and Nieminen 1990, Sipilä and Hyvärinen 1998, Amano et al. 2002, Olsen et al. 2025). A short postcanine tooth row has been reported to characterize the Saimaa ringed seal (Amano et al. 2002) and here we analyzed the dentition in greater detail. In the seals, the spacing of teeth, hence the length of the tooth row increases as the individual grows. Thus, as with other cranial and skeletal features, age-controlled sampling must be used for tooth row length. In contrast, individual tooth measurements are affected only by wear, rarely an issue with ringed seals as they lack exact occlusion and do not chew their food. Instead, the laterally compressed postcanines are used for biting and piercing of food, or retention of food in the oral cavity when the water is expelled (Loughlin 1982, Adam and Berta 2002, Hocking et al. 2017, Watanabe et al. 2020).

First, we compared the sizes of the largest four lower postcanines P_2_, P_3_, P_4_, and P_5_. These teeth are characteristic to species and in ringed seals have typically three to five sharp cusps. The postcanine lengths of a 326 specimen sample covering Arctic, Okhotsk, Baltic, Ladoga, and Saimaa show that the P_2_, to P_4_, fit between the longer Baltic, Ladoga, and Arctic, and the shorter Okhotsk specimens (Figure 4A). However, the last postacanine P_5_ (that is actually the first molar) is distinctly short in Saimaa, even shorter than the P_5_ of Okhotsk seals, the smallest of the studied ringed seals (*P* = 0.0000 to 0.0011, Figure 4A). Examining Saimaa tooth heights reveals that especially their anterior P_2_ and P_3_ are tall, and their overall height/length ratios, measuring relative height of the crown profile, are by far the highest among the populations (Figure 4B). Furthermore, the top-cusp angles, measuring relative height of the lateral cusps, are the smallest in Saimaa (Figure 4C, Table 2), further indicating a sharper crown profiles for the Saimaa ringed seal. This morphological pattern contrast with that of the characteristic ringed seal morphology observed in the other population samples, irrespective of their tooth size.

**Figure 4.**
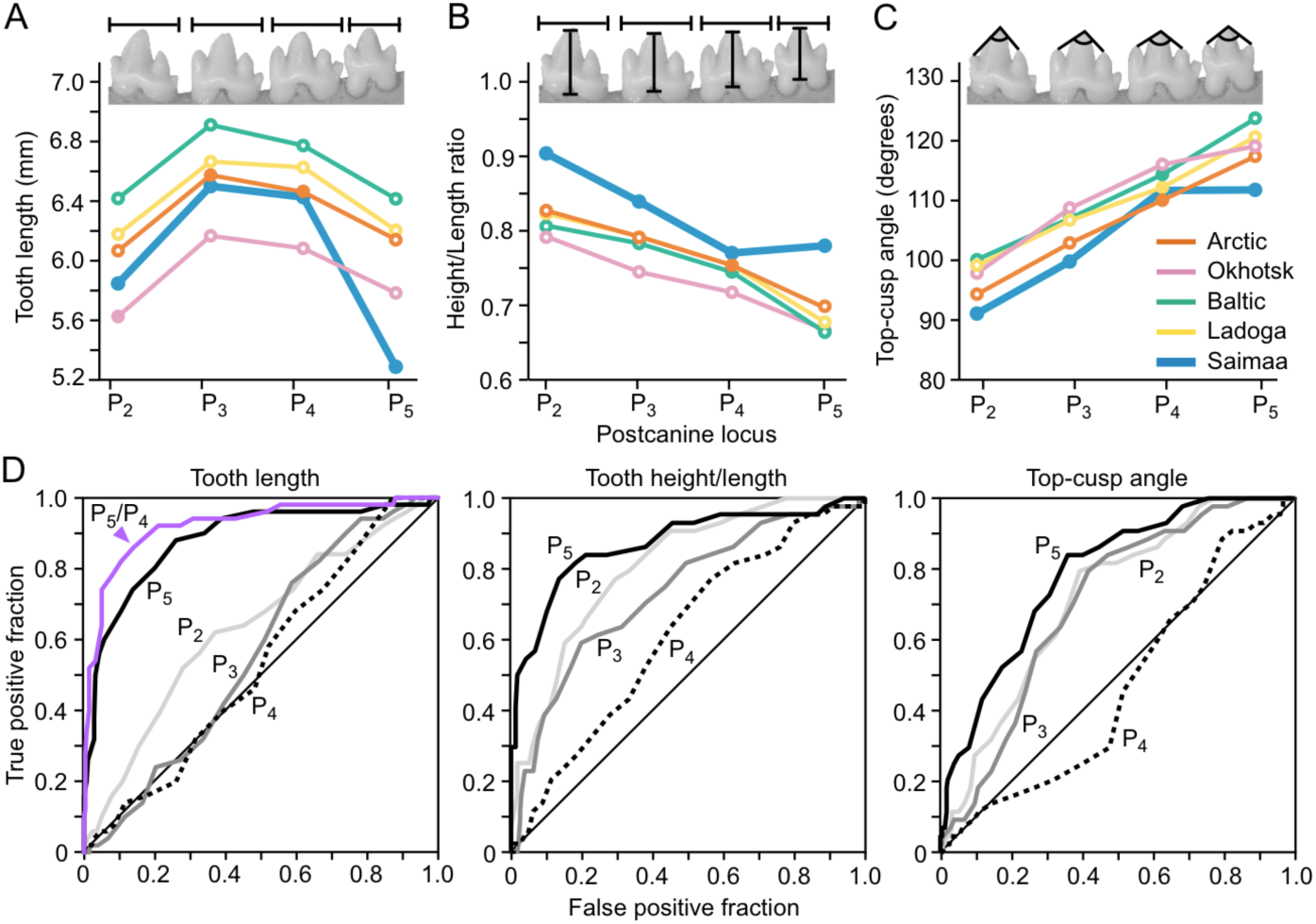
Saimaa ringed seals are distinct in their dental anatomy. (A) Mean lengths of lower postcanines P_2_ to P_4_ of Saimaa ringed seals are generally smaller than in Baltic and Ladoga, but only P_5_ is distinctly small compared to all the other populations. In the Saimaa dentition, the relative tooth heights (B), and the top-cusp angles measuring relative height of the lateral cusps (C), are the highest and smallest, respectively. These indicate shaper crown profiles in the Saimaa ringed seal, whereas the other populations retain the more characteristic ringed seal morphology irrespective of the size (A). Means differing from Saimaa with *P*-values below 0.05 are marked with open symbols in (A) to (C). For the *P* values, see Table 2. (D) ROC curves showing the performance of different teeth and measures in the classification of Saimaa ringed seals, P_5_ and to a lesser degree P_2_ performing the best for all the measures. For the length, removing absolute size variation by calculating P_5_/P_4_, improves the ROC curve further (purple).

**Table 2.** Permutation tests between Saimaa and other samples for tooth length, tooth height, height/length ratio (H/L), and top-cusp angle. *P*-values are one-tailed and except for the ones marked with asterisk, denote lower values for Saimaa lengths and top-cusp angles, and higher values for heights and height/length ratios. Tests are between group means using 10 000 permutations, *P*-values below 0.05 are marked in bold.

| Length | Ladoga | Baltic | Arctic | Okhotsk | Angle | Ladoga | Baltic | Arctic | Okhotsk |
| --- | --- | --- | --- | --- | --- | --- | --- | --- | --- |
| P2 | **0.0000** | **0.0000** | **0.0061** | 0.0623* | P2 | **0.0000** | **0.0000** | **0.0020** | **0.0051** |
| P3 | **0.0202** | **0.0001** | 0.2003 | **0.0019\*** | P3 | **0.0000** | **0.0001** | **0.0031** | **0.0000** |
| P4 | **0.0095** | **0.0002** | 0.3261 | **0.0018\*** | P4 | 0.2596 | 0.0641 | 0.0671* | **0.0260** |
| P5 | **0.0000** | **0.0000** | **0.0000** | **0.0011** | P5 | **0.0000** | **0.0000** | **0.0000** | **0.0006** |
| Height | Ladoga | Baltic | Arctic | Okhotsk | H/L | Ladoga | Baltic | Arctic | Okhotsk |
| P2 | **0.0145** | 0.1325 | **0.0026** | **0.0000** | P2 | **0.0000** | **0.0000** | **0.0000** | **0.0000** |
| P3 | **0.0126** | 0.3291 | **0.0020** | **0.0000** | P3 | **0.0001** | **0.0008** | **0.0000** | **0.0001** |
| P4 | 0.1663* | 0.1151* | 0.2045 | **0.0000** | P4 | 0.0571 | 0.1181 | 0.0513 | **0.0035** |
| P5 | 0.0559* | 0.0772* | **0.0012\*** | **0.0027** | P5 | **0.0000** | **0.0000** | **0.0000** | **0.0000** |

To examine the diagnostic value of tooth measures in identifying Saimaa ringed seals from the whole sample, we computed the receiver operating characteristic (ROC) curves for each tooth (Figure 4D). For the tooth length the areas under the ROC curves, which measure the performance, are 0.64, 0.56, 0.53, and 0.88 for P_2_, P_3_, P_4_, and P_5_, respectively (Figure 4D). Because the P_4_ appears to perform closest to random classification (area is closest to 0.5 and the line follows the diagonal in Figure 4D), we tested whether it could be used to remove overall size variation from P_5_ by calculating the P_5_/P_4_ ratio for each specimen. The P_5_/P_4_ ROC curve shows even better performance than the P_5_ length alone, enclosing 0.92 of the area (Figure 4D). A cut-off of approximately 0.88 for the P_5_/P_4_ ratio is indicated by the ROC curve as a diagnostic threshold value in the identification of Saimaa ringed seal material. Importantly, this size ratio is not affected by the overall small size of Okhotsk seals, and can also be used irrespective of age and sex, although females show better separation (Figure S2). In addition to tooth length, both height/length ratios and top-cusp angles of P_5_ provide diagnostic value, enclosing 0.87 and 0.79 of the ROC area, respectively (Figure 4D).

Because many seals swallow food with little or no processing, their tongue plays an important role in feeding, especially in species with a suction feeding strategy (Adam and Berta 2002). Although the availability of soft tissue anatomy is generally limited, next we examined the overall tongue morphology as it may help to explain the distinctive dental morphology of Saimaa ringed seals (Figure 5A). We observed that the shape of the tip of the tongue differs between the Saimaa and Baltic ringed seals, and only the former shows intermolar elevations on the lateral sides of the tongue (asterisks in Figure 5A). The width of the tongue is already over 70% of the final width at 20% from the tongue tip in the Saimaa specimens, whereas the Baltic specimen has much pointier shape, reaching 70% of the width closer to 40% from the tongue tip (Figure 5B). The tongue anatomy was recently examined in Antarctic seals and the crabeater (or the krill-eater seal, *Lobodon carcinophaga*) and the leopard seal (*Hydrurga leptonyx*) of that study are of special interest in comparison to ringed seals (Loza et al. 2023). The former is a suction and filter-feeding specialist that uses its complex teeth to retain the krill in the oral cavity. Its overall tongue shape appears to be an intermediate between the Saimaa and Baltic ringed seals (Figure 5B). In contrast, the leopard seal tongue is quite similar in overall shape to that of the Baltic ringed seal (Figure 5B). Whereas leopard seals consume krill much the same fashion as crabeater seals, they diversify their diet to vertebrate prey as they grow larger, switching from mainly suction-filter feeding to a grip-and-tear feeding mode.

**Figure 5.**
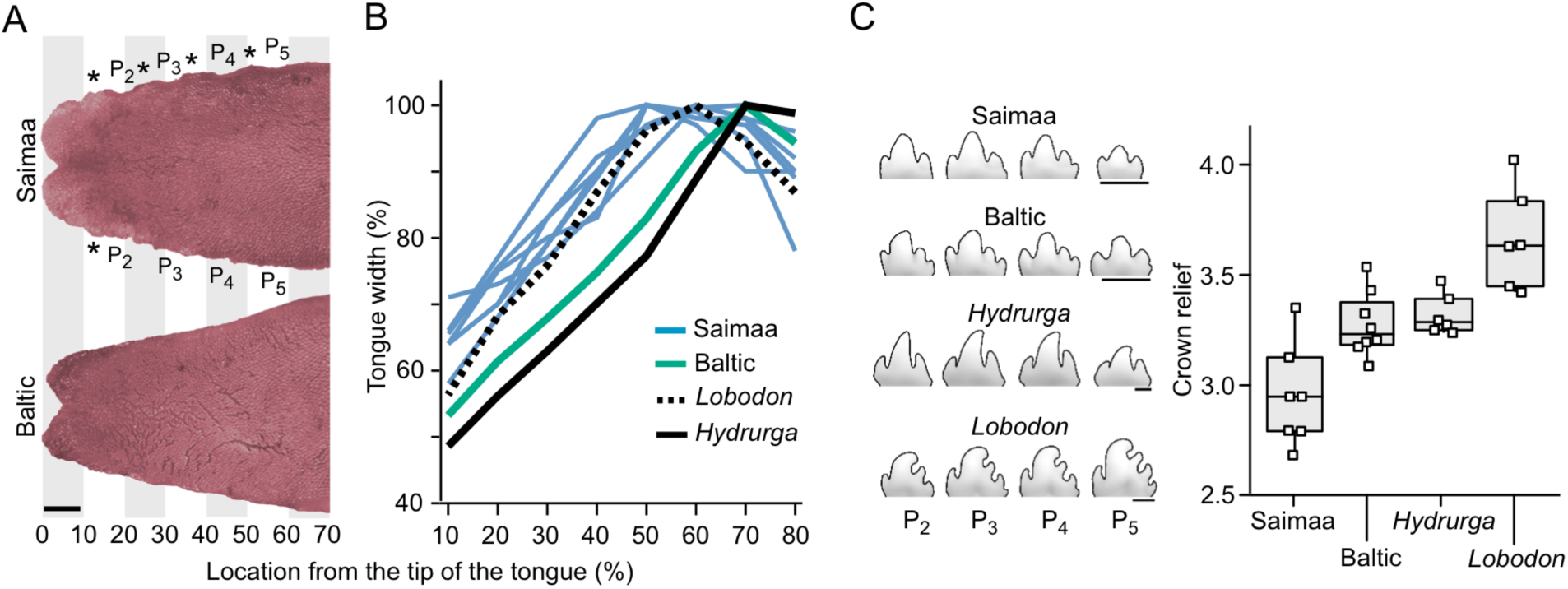
Tongue and dental morphology suggest increased suction but decreased filter-feeding niche for the Saimaa ringed seal. (A) Saimaa ringed seals have a broader tongue with a more rounded bifurcated tip, and intermolar elevations on the lateral side of the tongue (asterisks). (B) Comparison of tongue shapes with crabeater seal (*Lobodon*) and leopard seal (*Hydrurga*) tongues (from Loza et al. 2023) shows that the Saimaa ringed seal have more evenly broad tongue. (C) The cusp relief, measuring the filtering capacity of teeth is lower in Saimaa than in Baltic ringed seals, which in turn have comparable relief to that of the leopard seal. *P*-values between Saimaa sample and the other three are from 0.001 to 0.008 (one-tailed Mann-Whitney *U* test). To illustrate the tongue shapes, colors in (A) were adjusted to be similar. Scale bars for (A) and (C), 5 mm.

The known diets of ringed seals appear to incipiently align within the extremes of larger bodied crabeater and leopard seal diets. Ringed seals in the Arctic feed on small fish, typically around 10 cm, and rarely over 20 cm in length (Holst et al. 2001). However, Arctic ringed seals also consume zooplankton as seals in some regions show a seasonal switch to feeding on amphipod crustaceans, such as *Parathemisto libellula* (Lowry et al. 1980; Labansen et al. 2011, see also Geptner et al. 1988). In addition to seasonal feeding, amphipods are also regionally important dietary source for young seals (Holst et al. 2001). Although data is more limited for the Okhotsk population, also these seals have been reported to eat planktonic crustaceans in addition to fish (Fedoseev 1964, Geptner et al. 1988). Apart from the species composition, the Baltic ringed seals have comparable diets with the Arctic (Suuronen and Lehtonen 2012). In contrast, the Lake Saimaa is depauperate of large planktonic crustaceans and its seals feed almost entirely on small fish, such as vendace (*Coregonus albula*), smelt (*Osmerus eperlanus*) and perch (*Perca fluviatilis*) (Kunnasranta et al. 1999, Auttila et al. 2015).

Taken together, the addition of zooplankton to fish diet and the postcanines with well-developed cuspal comb (Ishihara et al. 2024) in marine ringed seals indicate adaptations to the filter feeding of zooplankton reminiscent of *Lobodon* and *Hydrurga* (Figure 5C). In comparison, cusp relief needed for filtering small prey is reduced in Saimaa ringed seals (Figure 5C, Figure 4). Rather, Saimaa ringed seals have sharper crown profiles emphasizing the central cusp at the expense of the lateral cusps, especially in the anterior postcanines (Figure 4). These morphological features suggest increased specialization to catching, biting and piercing of fish prey. This dental change is unlikely to be the result of a recent drift due to small population size because the broadened tongue implies integrated changes in the feeding apparatus. Kinematic analysis of feeding of marine ringed seals has shown them to be able to suction feed despite the lack of apparent specializations to suction in the skull morphology (Kienle et al. 2018). Considering the oral anatomy, we postulate that Saimaa ringed seals should have relatively strong suction feeding performance. Additional changes linked to feeding are suggested by subtle differences between jaw musculature between Saimaa and Baltic ringed seals (Laakkonen and Jernvall 2020), and the length of the intestinal tract of the Saimaa ringed seal being relatively shorter than those of marine ringed seals (Kunnasranta et al. 1999). Interestingly, the Ladoga ringed seals have not been documented to feed on zooplankton (Tormosov and Filatov 1979, but see Geptner et al. 1988), yet their teeth retain the typical ringed seal morphology (Figure 4). This suggests that the derived oral anatomy of the Saimaa ringed seal reflects a long regime of natural selection. To this end, the deep evolutionary roots of the Saimaa lineage have led to an opposite outcome compared to another long-isolated landlocked seal, the Baikal seal (*Pusa sibirica*). These seals have highly complex teeth, well suited for complementing their fish diet with suction-filter feeding of Lake Baikal’s abundant zooplankton (Watanabe et al. 2004, 2020, Ishihara et al. 2024).

## Discussion

Genome wide analyses of species relationships are providing more detailed, and increasingly more complex view into the history of many mammalian lineages (e.g., Cahill et al. 2018, Ferreira et al. 2021, Meneganzin and Bernardi 2023, Bertola et al. 2024). This can complicate taxonomic certainty in delineating species, and approaches integrating multiple lines of evidence have been advocated (Groves et al. 2016, Kitchener et al. 2022). For example, the traffic-light system proposed by Kitchener et al. (2022) uses three types of independent evidence: morphological, genetic and biogeographical. Accordingly, a simple genetic distance alone, even if large, does not provide taxonomic certainty. Integrative analyses combining multiple lines of evidence have shown that they can result in a reduction of the number of recognized species (e.g., van Elst et al. 2024), but also in identification of new taxa (e.g. Nater et al. 2017, Carroll et al. 2021). Given that the Lake Saimaa has formed a geographic barrier for dispersal during the last 10 kya, here we focused on genetic and morphological evidence to examine whether the ancestry of the Saimaa ringed seals could be older than the Lake Saimaa.

Our genetic analyses confirm that the seals in Lake Saimaa are not only clearly different from other ringed seals, but importantly, also have an independent and deeper evolutionary history. The Saimaa lineage has twice as distant split times from the three Atlantic ringed seals (Arctic, Baltic, Ladoga) than those have amongst them. Similarly, the Saimaa population has the greatest fraction of unique SNPs among the ringed seal populations, and the PCA component separating the Saimaa individuals from all the ringed seals explains nearly three times the amount of variance that the second component separating the four remaining ringed seal subspecies. Additionally, our analyses reveal a brief period of more recent contact between the Saimaa lineage and the lineages forming the current Ladoga and Baltic populations around 8–10 kya. The existence of this kind of intermittent gene flow is a recurrent observation of genetic admixture and introgression among many *bona fide* species in the wild (Huerta-Sanchez et al. 2014, Cahill et al. 2018, Ferreira et al. 2021, Bertola et al. 2024).

The deep genetic origin of the Saimaa lineage is further underscored by the morphological data. The distinct skull morphology of Saimaa ringed seals has been long recognized (e.g. Amano et al 2002), and here we demonstrate that their dental and tongue morphologies are indicative of specialization on feeding exclusively on fish. This specialized feeding niche differs from the other ringed seals that have more complex dental morphologies reflecting the ability to include zooplankton in their diet. Taken together, given the combination of genomic and phenotypic evidence, we propose that the Saimaa ringed seal is evolutionary more unique than previously thought.

The genetic differences and the timing of the population splits (Table 1, Figure 3C) are consistent with the scenario where the ancestral Saimaa population was formed from ice-dammed marine ringed seals in an eastern or southeastern refugium or possibly in multiple refugia latest during the Middle Weichselian, 90–50 kya ago, when large glacial lakes existed in the West Siberian Plains and White Sea Basin with water routes to the eastern edge of the Fennoscandian ice sheet (Mangerud et al. 2001, 2004, Krinner et al. 2004). We note that although ancient, as a landlocked marine population derived from eastern proglacial refugia, Saimaa seal is not unique. Relict marine fish and invertebrate species are found in modern Lake Saimaa and eastern lakes around the Baltic Sea basin (Savolainen 1975, Segerstrale 1976, Väinölä et al. 1994, 2001, Audzijonyte and Väinölä 2005, Säisä et al. 2005). It is conceivable that many of the other relicts may also have more ancient origins than previously thought, harboring unique evolutionary history (e.g. Rosing-Asvid et al. 2023).

Although Nordqvist (1899) originally described the Saimaa and Ladoga ringed seals, he also advocated for the extermination of both populations, due to their claimed nuisance to local fisheries (Nordqvist 1892). His goal of eradicating the Saimaa seal was nearly realized, with fewer than 100 individuals remaining when the population finally gained legal protection in 1955 (Kunnasranta et al. 2021). The population has gradually recovered since then, reaching almost 500 individuals in 2024 (https://www.metsa.fi/en/nature-and-heritage/species/saimaa-ringed-seal/). As Saimaa seals depend on snow lairs for birthing and nursing, ongoing anthropogenic climate change presents a significant long-term threat (Jakkila et al. 2024). However, the Saimaa seal has survived the previous post-glacial warm period and is likely able to do so in the future, if given the chance.

## Materials and methods

### Material examined

DNA samples from ringed seals were obtained from 12 unique sampling localities in the northern hemisphere, greatly improving the coverage in the eastern regions compared to the previous analyses [Figure 1, distribution ranges drawn after Kelly et al. (2010)]. The sample sources are described in Löytynoja et al. (2023) and Rosing-Asvid et al. (2023). The sources of the new samples for Baltic and Ladoga are the same as in (Löytynoja et al. 2023). Alaska samples are from Prudhoe Bay (001B98, AF7110), Peard Bay (G003), and Bering Sea TV13-15); Chukchi Sea samples are from Lorino; Okhotsk Sea samples are mainly from Chkalov Island and Odyan Bay (TV16-18); Pechora Sea samples are from Vaigach Island; and White Sea samples from Belomorsk archipelago. Samples are from bycaught seals or dead beachcast animals collected for research. Based on earlier analyses (Löytynoja et al. 2023, Rosing-Asvid et al. 2023), the whole-genome-sequenced individuals were ranked by quality (maximizing the sequencing coverage and data completeness) and at most five individuals from each population were included for full genome analyses (in total 46 samples). Four spotted seals (*Phoca largha*; accession numbers SAMN08238620, SAMN16895771, SAMN31577600 and SAMN31577601; Park et al. 2018, Dudchenko et al. 2017, Xu et al. 2024) were included for SNP polarization. The mitochondrial DNA (mtDNA) was analyzed separately and 162 unique ringed seal haplotypes and the four spotted seal genomes were included. For the dental data, Saimaa material are from the University of Eastern Finland (UEF) and Metsähallitus (the Finnish Forest Administration), Ladoga material from Finnish Museum of Natural History (FMNH), University of Helsinki, University of Eastern Finland and Metsähallitus, Baltic material from Finnish Museum of Natural History (FMNH), Arctic Greenland material (East and West Greenland, including Canadian Arctic Archipelago) from National History Museum of Denmark (NHMD), Arctic Alaska material (Beaufort, Chukchi, and Bering Sea) from University of Alaska Museum of the North (UAM), and Arctic Kara Sea and Sea of Okhotsk material from the National Museum of Nature and Science of Tokyo (NSMT). The four distal postcanines show the largest range of variation among phocid species, and their intra- and interspecies variation have been studied extensively previously (Jernvall 2000, Savriama et al. 2018, Ishihara et al. 2024). Additionally, we investigated soft tissue anatomy of tongues from freshly defrozen seal cadavers (*N*=8, from Metsähallitus (the Finnish Forest Administration). The type specimens are presented in a separate chapter. Comparative dental specimens to calculate cusp relief originated from the collections of the Finnish Museum of Natural History (FMNH), the Swedish Museum of Natural History (NRM), and the National Museum of Victoria, Melbourne (NMV).

### Genomic analyses

#### Data mapping and nuclear genome variant calling

The DNA sequencing was performed on Illumina NextSeq500 and NovaSeq 6000 platforms (Löytynoja et al. 2023, Sundell et al. 2023). Read data for ringed seal samples and four spotted seal samples were aligned as described previously (Olkkonen and Löytynoja 2023). For the selected 46 ringed seals and four spotted seals, variants were called by applying GATK4 (v.4.2.5) (Poplin et al. 2018) HaplotypeCaller for each sample, joining the data with CombineGVCFs, and applying GenotypeGVCFs to the joint data. The sequencing depth of the studied ringed seal samples varied from 5.0x to 29.5x with the mean coverage of 10.6x, and of spotted seal samples from 34.8x to 115.7x with the mean coverage of 55.7x.

#### Mitogenome analyses

Variants within the mitochondrial genome was called from all sequenced individuals applying the same GATK4 tools but defining the ploidy 1. Protein-coding regions were inferred by mapping the peptide sequences of the *Pusa hispida* mitochondrion (NC_008433) against the reference mtDNA contig with miniprot (Li, 2023). Binary SNP variants, totalling 1,434, within the CDS regions were extracted and samples with missing data were discarded. The VCF data were converted to FASTA sequences using the vcf-to-tab tool from the VCFtools package (Danecek et al. 2011) and the script from https://github.com/JinfengChen/vcf-tab-to-fasta modified to work on haploid data. Identical haplotypes (all within a subspecies) were removed, leaving 162 ringed seals and four spotted seals as the outgoup. The phylogenetic tree was inferred with RAxML v.8.2.12 (Stamatakis, 2014) using the model ASC_GTRGAMMA and correcting for the invariable sites (totalling 9,945) missing from the VCF file with the option -- asc-corr=felsenstein. For the search, the rapid option combining the ML search (20 replicates) and the bootstrap analysis (1000 replicates) was selected (-f a -# 1000). The resulting tree was visualised in R using the ggtree package (Yu et al. 2017).

#### Data filtering and SNP analyses

The binary SNP data for the full nuclear genomes were filtered and thinned using BCFtools (v.1.9) (Danecek et al. 2021) and Plink (v.1.9) (Chang et al. 2015). Sites falling outside the positively masked regions or inside the repeat-masked regions (Olkkonen and Löytynoja 2023), showing no variation among ringed seals or having >10% of missing data were excluded. The remaining data, totalling 25,697,406 sites, were further thinned by requiring sites to be at least 1 kbp apart, leaving 1,807,907 variant sites.

A principal component analysis was performed on the thinned data (1.8M sites) with smartpca (v.16000)(Price, et al. 2006) and the results were visualized with R (R-Core-Team 2020) using the ggplot2 (Wickham 2016) package. In smartpca, the normalization option (usenorm) was turned off and the analysis was performed for the full 46 individuals set and for an even sampling of five individuals per subspecies (Elhaik, 2022), selected based on data completeness and, in the case of the Arctic subspecies, picking individuals from different geographic locations (Alaska, Svalbard, White Sea, Pechora Sea, Chukchi Sea). Starting from the 1.8M sites data, allele-sharing among different subspecies was studied using the same five individuals per subspecies, including the four spotted seals for derived allele inference. With samples varying from one to five individuals, the presence of a derived allele in each ringed seal subspecies was recorded, and the proportion of derived allele patterns unique to the specific subspecies was calculated. The statistic is highly dependent on the MAF limit applied. With full data (MAF=0), a large proportion of unique alleles are singletons and reflect the recent demographic events and the current effective population size. We applied MAF=0.05 (requiring at least five derived alleles among the 92 sampled genomes in the full data) to understand the more distant demographic events; however, one cannot rule out the founder effect and recent genetic drift inflating the Saimaa statistics. The proportion of private variants is expected to decrease with the number of individuals sampled and, in theory, plateau and stop when all individuals are included. The allele frequencies in each subset were computed with VCFtools and processed with bash and R scripts.

#### Demographic analyses

Demographic analyses were performed with MSMC-IM (Wang et al. 2020) using the whole-genome data from Löytynoja et al. (2023). For within-population analyses, ten two-individual sets (i.e., four chromosomes) were randomly chosen from each population and their coalescent rate histories were computed as explained in Löytynoja et al. (2023). For cross-population coalescence rate analyses, six pairs of individuals for each population combination (i.e., two chromosomes from each population, two populations per analysis) were randomly chosen and their coalescent rate histories were computed using the default time segmenting. The combined rate estimates were analysed using MSMC_IM.py with the recommended parameters and the mutation rate (*μ*) 1.826e-8 estimated for polar bear (Liu et al. 2014). We consider this a robust estimate as it falls within the mutation rates estimated for different seals (Peart et al. 2020), and using different estimates does not affect the overall pattern of results. The summary results were computed and visualized with R using the ggplot2 package and assuming a generation time (*g*) of 10 years.

#### Phenotypic analyses

Lower tooth rows were photographed from the lingual side and maximum tooth lengths, maximum heights, top-cusp angles, and cusps numbers were tabulated using Fiji (Schindelin et al. 2012) and additional analyses were done with PAST (Hammer et al. 2001). Although the dentitions of phocids show high degree of intraspecies variation, the seals are born with their permanent tooth crowns fully formed and in the process of erupting into the oral cavity. Therefore, in contrast to other skeletal features, dental measures do not change after birth. We report the analyses for the four, right side postcanines (P_2_, P_3_, P_4_, P_5_), and only for individual specimens from which the measures could be obtained for all the four teeth. We excluded specimens with dental anomalies such as supernumerous teeth. Cusp number and tooth length were measured for 326 individuals and crown height and top-cusp angle for 281 individuals due to cracked or slightly worn specimens. Height and angle measurements were estimated for cases where only the enamel cap of the cusp tip was missing. Exclusion of these data did not alter the results. Cusp were tabulated regardless how small they are. Relative cusp height was measured using the top-cusp angle (Jernvall 2000, Salazar-Ciudad and Jernvall 2010) and the cusp relief as the length of the crown perimeter divided by the length of the tooth at the crown base. *P*-values in Figure 3 and Table 2 are one-tailed and obtained between group means using 10 000 permutations. The receiver operating characteristic (ROC) curves were used to obtain correct classification probabilities. ROC curves have been more typically used for medical data analyses, but can also be applied for taxonomical questions (e.g., Dos Santos et al. 2011). For tooth lengths and top-cusp angles, smaller values denote true positive. Because seal tongue plays an important role in feeding, especially in species with a suction feeding strategy (Hocking et al. 2017, Loza, et al. 2023), we compared the overall morphology of Saimaa ringed seal tongue with that of phocids from the literature (Loza, et al. 2023). Tongues from cadavers were photographed in defrosted stage and compared to those in the literature. To obtain a robust measure of shape apart from size and variation caused by differential preservation of soft tissue anatomy, were divided tongue images into ten-percent bins from the tip to the end of the posterior body, and tabulated widths as relative to the maximum width of the tongue.

## Data availability

Most of the genome data have been previously used and are available at the European Nucletiode Archive (ENA) under accession numbers PRJEB56329 and PRJEB56317. The newly generated 16 whole-genome sequencing datasets are deposited under PRJEB56317 with accession numbers ERS23016468-ERS23016483. Instructions for replicating the computational analyses are provided at https://github.com/ariloytynoja/pusa_saimensis

## Acknowledgements

For tissue samples, we thank the University of Eastern Finland and Metsähallitus (Saimaa); M. Verevkin (Ladoga); P. Timonen, S. Oksanen, and J. Vierimaa (Baltic); D. Tallmon, A. Rosing-Asvid, S. Ferguson, R. Dietz, M. Zasypkin, P. Strelkov, N. Medvedev and particularly O. Shpak (Arctic). For access to dental material, we thank the curators of Finnish Museum of Natural History, National History Museum of Denmark, the National Museum of Victoria (Melbourne), the Swedish Museum of Natural History, and University of Alaska Museum of the North (Fairbanks). For advice on taxonomy and analyses we thank S. Viranta, K. Leppälä, and X.-Y. Feng. We thank the personnel of the DNA sequencing and genomics laboratory for performing the NGS samples preparation and sequencing, and the CSC – IT Center for Science for the computational resources. The Jane and Aatos Erkko Foundation (JJ and PA), the LIFE Programme of the European Commission (PA, LIFE19NAT/FI/000832), Research Council of Finland (JJ) are acknowledged for their financial support.

## Supporting figures

**Figure S1.**
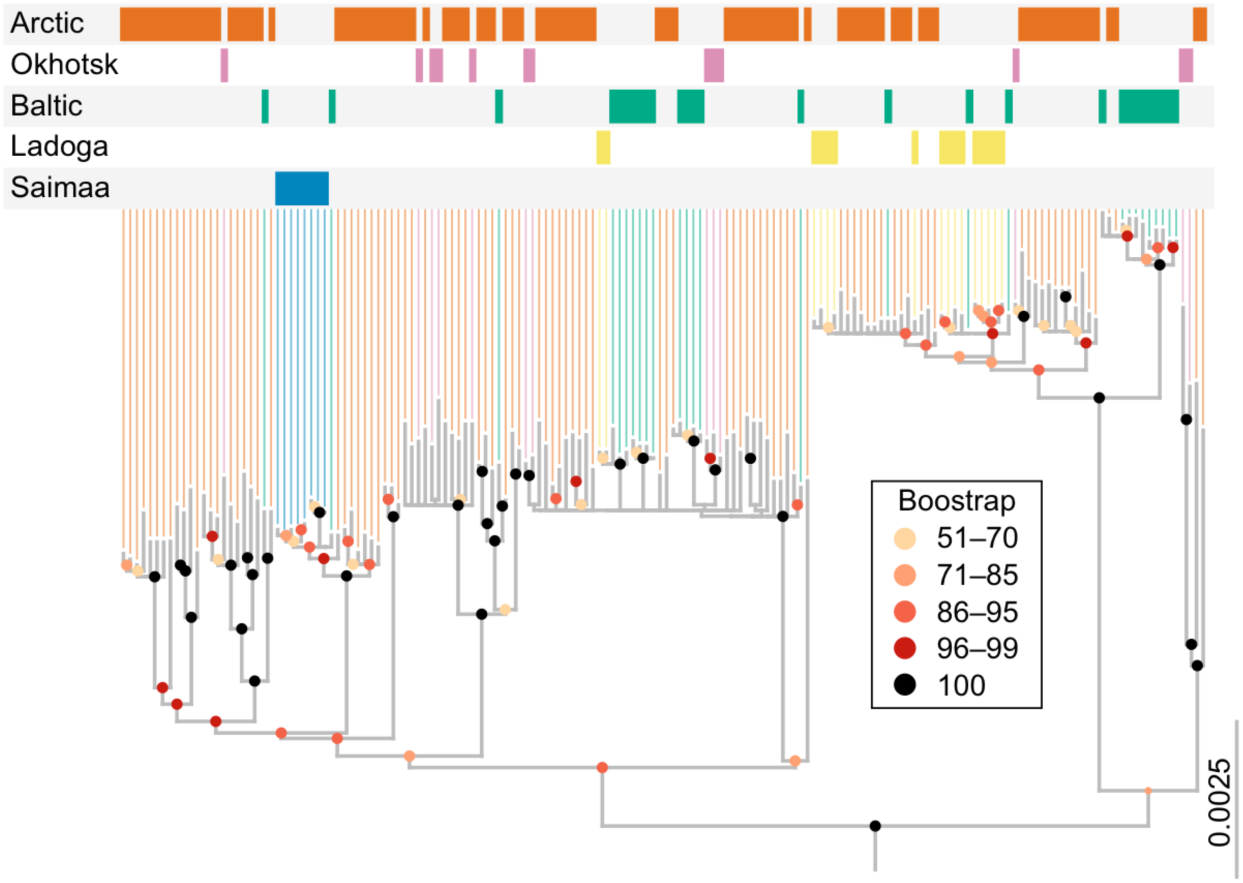
The maximum likelihood tree for the unique mitogenome haplotypes rooted with four spotted seals. The dots at internal nodes indicate bootstrap support values greater than 50%. In contrast to the four ringed seal subspecies, all Saimaa samples (in blue) cluster tightly together and form an evolutionary lineage of their own.

**Figure S2.**
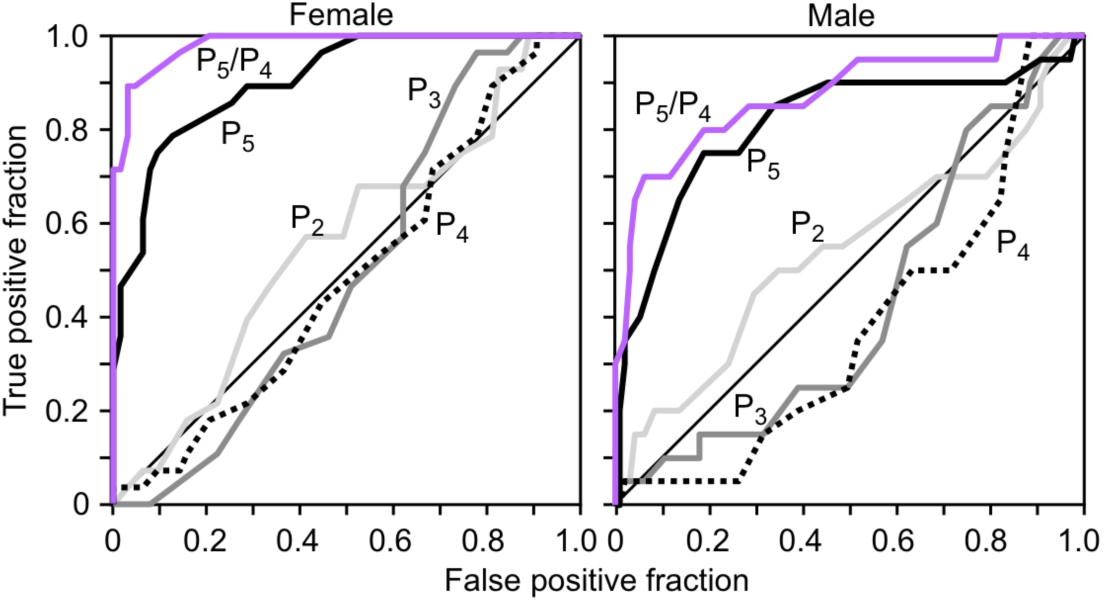
The receiver operating characteristic (ROC) curves for tooth lengths plotted separately for female and male specimens. Of the 326 specimens measured, 96 (28 from Saimaa) and 115 (20 from Saimaa) are assigned to female and male, respectively. For both sexes, the P_5_/P_4_ performs the best, but female seals show almost perfect classification. Areas below the P_5_ and P_5_/P_4_ curves are 0.90 and 0.98 for females, and 0.81 and 0.87 for males.

